# Corticolimbic structure-function coupling is sensitive to childhood adversity and buffers adversity-related symptoms during development

**DOI:** 10.64898/2025.12.22.696017

**Authors:** Lucinda M. Sisk, Taylor J. Keding, Allison Drew, Emilie Ma, Golia Shafiei, Matthew Cieslak, Theodore D. Satterthwaite, Dylan G. Gee

## Abstract

Childhood adversity is a potent predictor of mental health problems across the lifespan, and a rich cross-species literature implicates stress-sensitive corticolimbic circuits in adversity-related psychopathology. Structure-function coupling (SFC) is a promising multimodal marker that is sensitive to developmental plasticity. While emerging evidence suggests cortical SFC is sensitive to adversity exposure during early childhood, it is unknown how SFC in stress-sensitive corticolimbic circuits links adversity exposure with mental health across development. We examined associations between adversity exposure, transdiagnostic symptomatology, and both amygdala-cortical and hippocampal-cortical SFC across development in a large sample of youth (N = 607, 39% F). Results revealed that adversity exposure moderated age-related change in amygdala-vmPFC SFC (*p* = .007), such that youth exposed to higher, but not lower, levels of adversity showed an age-related increase in amygdala-vmPFC SFC. Further, amygdala-vmPFC SFC buffered the effect of adversity on internalizing symptoms (*p* = .012), such that those youth with higher adversity exposure who had stronger amygdala-vmPFC SFC also displayed lower internalizing symptoms. Separately, higher adversity exposure was associated with lower hippocampal-limbic network SFC (*p* = .037), which moderated the effect of adversity on internalizing symptoms (*p* = .010). These findings highlight that structural and functional neurodevelopment of amygdala-vmPFC and hippocampal-limbic network connections may adapt in distinct ways to support mental health following adversity, with implications for risk and resilience against internalizing psychopathology.

**Significance Statement:** Delineating how childhood adversity alters neurodevelopment is critical to understanding the origins of adversity-related risk for mental health disorders. Here, we leverage a multimodal marker of the correspondence between brain structure and function to test how corticolimbic circuits are jointly shaped by adversity exposure. Results reveal that the structure-function coupling of amygdala-vmPFC and hippocampal-limbic networks is associated with adversity exposure and differentially moderate risk for internalizing psychopathology. These findings suggest key roles for neural stress regulation and memory circuits in adapting to support optimal functioning following adversity exposure, and highlight the concordance of neural structure and function as an important indicator of individual-level risk and resilience.

## Introduction

Childhood adversity is a potent risk factor for mental health problems at the population level (Copeland et al., 2018; Felitti et al., 1998; Green et al., 2010), but predicting individual risk following adversity exposure remains challenging due to vast heterogeneity in outcomes (Meehan et al., 2022). Experience-dependent neurodevelopmental changes following adversity exposure may contribute to this variability by inducing changes in brain structure and function (Andersen, 2003; Greenough et al., 1987; McLaughlin & Gabard-Durnam, 2019). Cross-species studies have established that adversity exposure is associated with altered development of white matter microstructure and functional connectivity, particularly among stress-sensitive regions such as the hippocampus, amygdala, and ventromedial prefrontal cortex (vmPFC) (Eiland et al., 2012; McEwen et al., 2016; Mitra et al., 2005; Tottenham & Sheridan, 2010). Here, we examine how the coupling between structural and functional connectivity in key corticolimbic circuits develops and relates to adversity exposure and psychopathology.

Amygdala-prefrontal and hippocampal-prefrontal connections play a critical role across species in processing threat-related and emotional information (e.g., Casey et al., 2019; Tottenham & Gabard-Durnam, 2017). Individuals exposed to adversity show altered functional (Herringa et al., 2016; McLaughlin et al., 2019; Sisk et al., 2025; Tottenham et al., 2011; van Harmelen et al., 2013) and structural (Eluvathingal et al., 2006; Hanson et al., 2015; McCarthy-Jones et al., 2018) neurodevelopment in these circuits, potentially underpinning differences in cognitive and emotional functioning. Altered neurodevelopmental timing may be one mechanism through which such differences emerge (e.g., Belsky, 2019; Callaghan & Tottenham, 2016; McLaughlin & Gabard-Durnam, 2022; Sisk & Gee, 2024). Indeed, accelerated corticolimbic neurodevelopment may reflect ontogenetic adaptation to a challenging environment (e.g., DeJoseph et al., 2024; Ellis et al., 2022) and can buffer against worsened mental health symptoms (Chan et al., 2024; Gee et al., 2013; Herzberg et al., 2021). However, despite substantial evidence for the effects of adversity on corticolimbic structure and function independently, it is unknown how adversity exposure may relate to structure-function coupling (SFC) within these circuits. As SFC reflects plasticity of developing circuits (Fotiadis et al., 2023) and can predict behavioral phenotypes above and beyond structure or function alone (Gu et al., 2021; Jiang et al., 2019), addressing this question has potential to advance understanding of heterogeneity in adversity-related outcomes.

White matter structure fundamentally constrains patterns of neural activity across levels of analysis (Shen et al., 2012), yet jointly evaluating these metrics has only recently become popularized through advances in computational power and network neuroscience (Bassett et al., 2018). SFC estimates how structural connectivity relates to functional connectivity at the connectome level (Baum et al., 2020; Fotiadis et al., 2024). SFC varies across the brain and changes with neurodevelopment—for instance, SFC is higher in unimodal sensorimotor cortex and lower in transmodal association cortex, reflecting increased myelin content and inhibitory neurotransmission in unimodal versus transmodal regions (Baum et al., 2020; Fotiadis et al., 2023; Gu et al., 2021). SFC is also sensitive to environmental input. Indeed, prenatal adversity exposure is associated with accelerated SFC development in association networks (Chan et al., 2024). However, the majority of SFC studies to date have focused solely on the cortex, omitting stress-sensitive corticolimbic circuits. Thus, how hippocampal-cortical and amygdala-cortical SFC changes across development remains poorly understood.

We addressed these gaps by examining SFC in hippocampal-cortical and amygdala-cortical circuits in a large sample of youth aged 5-18. We hypothesized that hippocampal-vmPFC and amygdala-vmPFC SFC would increase with age, and that adversity exposure would moderate age-related changes in SFC such that youth with higher levels of adversity exposure would show more mature patterns of SFC relative to youth with lower levels of adversity exposure. As accelerated corticolimbic circuit development may confer resilience against psychopathology in youth exposed to adversity, we hypothesized that SFC of hippocampal-vmPFC and amygdala-vmPFC connections would moderate associations between adversity exposure and transdiagnostic symptoms. Specifically, we expected that an accelerated pattern of development in youth with higher levels of adversity exposure would be associated with reduced symptom burden.

## Methods

### Participants

Participants were 607 youth ages 5-18 who participated in the Healthy Brain Network (HBN) study (Alexander et al., 2017). Participants were recruited through a community referral model that aimed to capture broad heterogeneity in development and psychopathology (Alexander et al., 2017). Exclusion criteria included acute safety concerns, behavioral or cognitive differences that would hinder study participation, and medical concerns that might confound neuroimaging research (e.g., neurodegenerative disorders) (Alexander et al., 2017). Prior to study participation, 18-year-old participants provided written informed consent, and a parent or legal guardian provided written informed consent for participants younger than 18. Participants and a parent attended a series of four sessions at one of three sites in the broader New York City area. Sessions included diagnostic interviews, child-report and parent-report questionnaires, and an MRI scanning session (Alexander et al., 2017). Detailed demographic information for participants included in the present study is presented in **Table 1**.

**Table 1.**
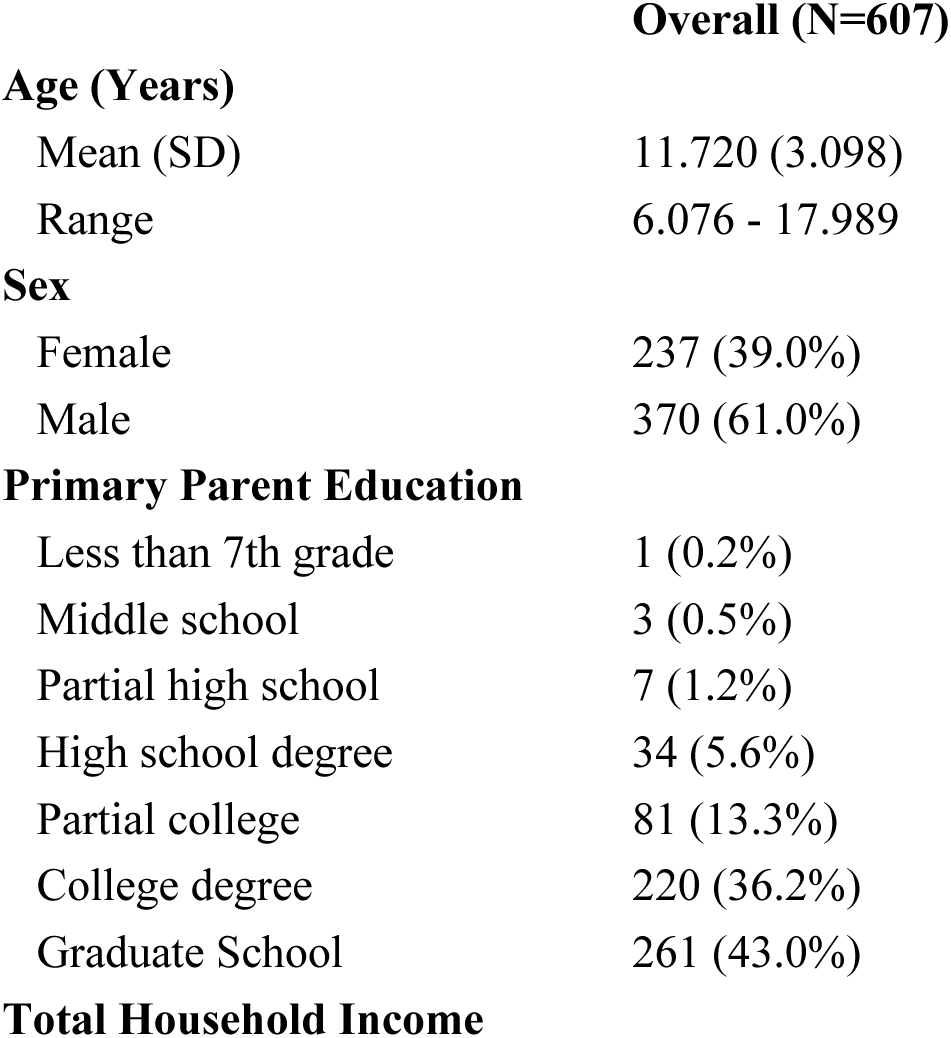
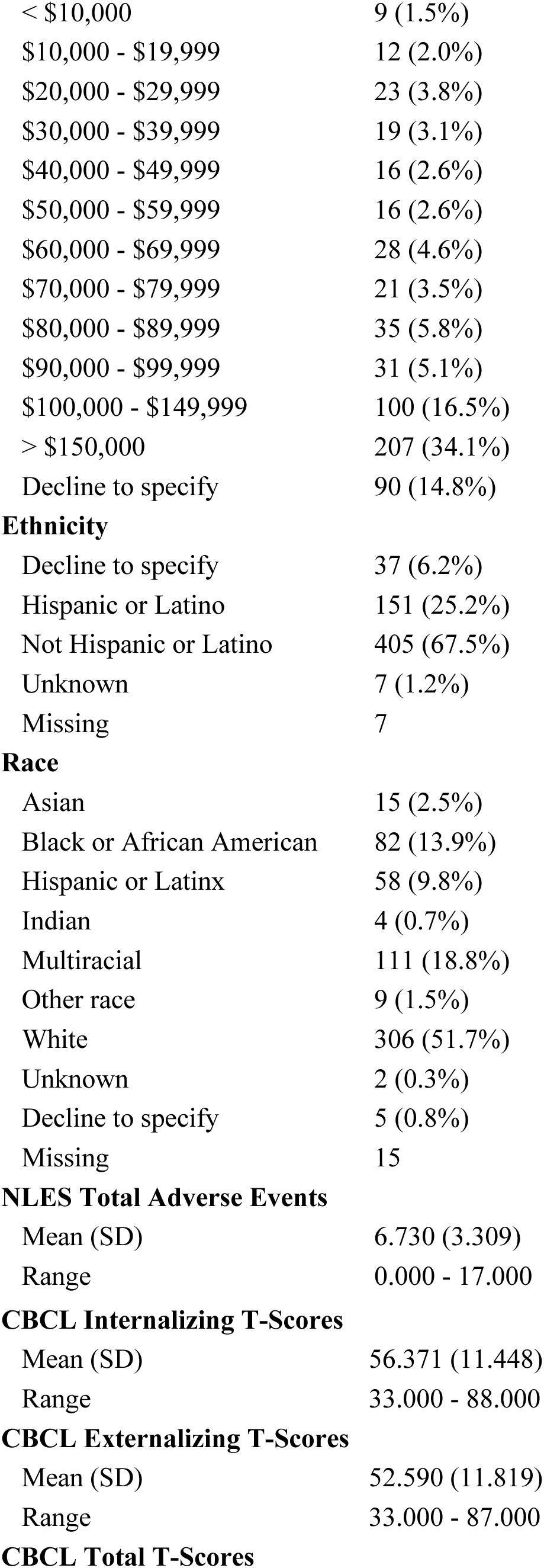
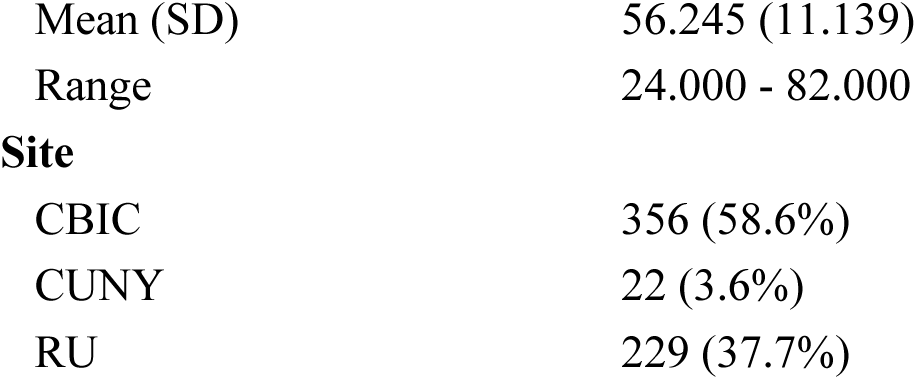
Demographics for the sample included in the present study. CBCL = Child Behavior Checklist; CBIC = Citigroup Biomedical Imaging Center. CUNY = City University of New York. NLES = Negative Life Events Scale. RUBIC = Rutgers University Brain Imaging Center.

### Adversity exposure

Participants’ exposure to adversity was queried using the Negative Life Events Scale (NLES) (Tiet et al., 2001). The NLES comprises 21 questions assessing life events such as experiencing severe injury or illness, having a parent lose a job, or moving schools. While HBN collects both parent-and child-reported NLES questionnaires, we chose to use parent-reported NLES scores in the present study to minimize missingness. We used the ‘total_events’ variable, which is a count of the total number of negative life events endorsed.

### Clinical measures

The Child Behavior Checklist (CBCL; Achenbach, 1999) is a widely used and well-validated measures that was used to assess clinical symptoms. CBCL scores from the HBN study were normally distributed and reflected a broad range of symptom severity experienced by participants in the HBN study (Alexander et al., 2017). Given our primary focus on adversity and neurodevelopment, as well as the fact that adversity is associated with transdiagnostic symptoms, we chose to examine T-scores representing internalizing, externalizing, and total symptoms, rather than disorder-specific subscales.

### MRI data acquisition

MRI data were acquired using Siemens 3-Tesla scanners; data collected using 1.5-Tesla scanners at the Staten Island site were excluded from this analysis. Data collected at the Rutgers University Brain Imaging Center used a Siemens Tim Trio scanner, and data collected at the CitiGroup Cornell Brain Imaging Center and the CUNY Advanced Science Research Center used Siemens Prisma scanners. Data were collected using a 32-channel head coil. For detailed information regarding the MRI acquisition parameters, see the original data descriptor paper (Alexander et al., 2017).

### Data access

Preprocessed diffusion-weighted imaging data from the HBN study were obtained through a publicly available repository (Richie-Halford et al., 2022). Subsequent reconstruction and connectome creation were performed using QSIRecon (Cieslak et al., 2021, 2025), as described below. Preprocessed resting-state timeseries data and processed anatomical images from the HBN study were obtained from the Reproducible Brain Charts (RBC) project (Shafiei et al., 2024, 2025). Subsequent parcellation and connectome construction were performed using FSL tools (Jenkinson et al., 2012) and custom python code (van Rossum, 1995).

### Quality assessment

Structural and functional imaging data underwent rigorous quality assessment. Structural diffusion data were assessed through a combination of expert raters, community raters who evaluated images through a web portal, and machine learning models trained on these ratings (Richie-Halford et al., 2022). In the present study, data with a quality score of 0.3 or above were included, and the quality score was included as a covariate in all neuroimaging analyses. Functional data image quality was assessed primarily through motion metrics. fMRI runs with framewise displacement (FD) values less than or equal to 0.2 and normalized cross-correlations greater than or equal to 0.8 were considered adequate for use (Shafiei et al., 2024, 2025). Data that did not meet these criteria were excluded from the present study. T1-weighted anatomical images were manually evaluated by 2-5 community raters who participated in “Swipes for Science” (Keshavan et al., 2019). Data were subsequently assigned a rating of “Pass”, “Artifact”, or “Fail” based upon these manual reviews. Only data rated as “Pass” were included in the present study.

### Anatomical data processing

T1-weighted anatomical images obtained through the RBC data release (Shafiei et al., 2025) were processed using FreeSurfer (Fischl, 2012) and sMRIPrep (Esteban et al., 2024), yielding commonly used measures of brain structure. FreeSurfer derivatives were included in the diffusion image reconstruction pipeline to improve anatomical accuracy.

### Diffusion connectome data

Diffusion data reconstruction was performed using QSIRecon version 1.0.0rc3 (Cieslak et al., 2021, 2025), which is based on *Nipype* 1.5.1 (Gorgolewski et al., 2011; Gorgolewski et al., 2018); RRID:SCR_002502. The following boilerplate descriptions from QSIRecon are reported in alignment with recommendations for best practices for reproducible neuroscience (Cieslak et al., 2021, 2025). Details on the QSIPrep pipeline used to preprocess these data can be found in the data release paper (Richie-Halford et al., 2022).

### MRtrix3 processing

FreeSurfer outputs were registered to the QSIPrep outputs to improve anatomical accuracy. Multi-tissue fiber response functions were estimated using the dhollander algorithm. Fiber Orientation Districutions (FODs) were estimated via constrained spherical deconvolution (Tournier et al., 2004; Tournier et al., 2008) using an unsupervised multi-tissue method (Dhollander et al., 2016, 2019). FODs were intensity-normalized using mtnormalize (Raffelt et al., 2017).

### Anatomical processing

T1w-based spatial normalization calculated during preprocessing was used to map atlases from template space into alignment with DWIs. Brain masks from antsBrainExtraction were used in all subsequent reconstruction steps. Cortical parcellations were mapped from template space to DWI data using the T1-weighted-based spatial normalization.

### Structural connectome construction

Diffusion data were parcellated using the 4S atlas (https://github.com/PennLINC/AtlasPack), which includes 400 cortical parcellations from the original Schaefer atlas in addition to 56 subcortical segmentations from atlases representing thalamic nuclei (Najdenovska et al., 2018), the cerebellum (King et al., 2019), the hippocampus and amygdala (Glasser et al., 2013), and additional subcortical nuclei such as the hypothalamus, caudate nucleus, and putamen (Pauli et al., 2018). Tractography was performed using tckgen, which uses the iFOD2 probabilistic tracking method to generate 1e7 streamlines with a maximum length of 250mm, minimum length of 30mm, FOD power of 0.33. Weights for each streamline were calculated using SIFT (Smith et al., 2013), an approach that improves the biological accuracy of the streamline count by using the underlying spherical deconvolution of the diffusion signal to weight streamlines, and were included while estimating the structural connectivity matrix. Structural connectomes were generated representing the streamline counts between all atlas regions, and resulting measures were scaled by the inverse of the node volumes in order to account for the inherently greater number of streamlines that emerge from larger relative to smaller atlas regions (Hagmann et al., 2008; Tournier et al., 2019).

### Structural connectome thresholding

To reduce the incidence of potentially spurious streamline connections present in the structural connectivity matrices, we applied a consistency-based thresholding approach (e.g., Baum et al., 2020; Gong et al., 2009; Li et al., 2012; Roberts et al., 2017) that retains streamlines that are consistently present across participants regardless of their relative strength. In line with previous work (Baum et al., 2020), we thresholded participants’ connectivity matrices at the 75th percentile for edge weight, removing the 25% most inconsistent connections across participants. We subsequently normalized each matrix at the subject level by dividing each edge by the total weight of network connections in order to facilitate comparisons across participants (Baum et al., 2020).

### Functional Connectome Data

Functional imaging data were processed using the Configurable Pipeline for the Analysis of Connectomes (C-PAC) (Li et al., 2024) which is based on fMRIPrep (Esteban et al., 2019). Specific details on functional data preprocessing can be found in the RBC data release paper (Shafiei et al., 2024, 2025).

### Functional connectome construction

As resting-state connectivity estimates stabilize with ∼10 minutes of data (Gonzalez-Castillo et al., 2014), we concatenated the two 5-minute preprocessed resting-state timeseries along the time dimension, which was then parcellated using the 456-node 4S atlas. Pearson correlations were computed between each node’s timeseries to generate connectivity matrices for each participant.

### Network extraction

#### Identifying vmPFC nodes

Given our a priori interest in corticolimbic neural architecture, we sought to robustly identify a subset of atlas regions representing the vmPFC. To this end, a vmPFC mask was created by merging all regions of the Mackey Ventromedial Prefrontal Cortical atlas (Mackey & Petrides, 2014) into a single region of interest (ROI). We then estimated the degree of overlap between parcels of the 4S atlas with the Mackey atlas ROI. 4S parcels where 50% or more of voxels overlapped with the Mackey atlas ROI were retained as a part of the ‘vmPFC’ network.

#### Subcortical-network measures

In order to isolate SFC between cortical and subcortical regions of interest, we extracted several subsets of nodes prior to computing SFC. Specifically, from the functional and structural connectomes, we separately extracted networks representing connectivity among bilateral hippocampal and amygdala segmentations and cortical regions, separately, in the visual, somatomotor, dorsal attention, salience, limbic, frontoparietal, and default mode networks according to Yeo 7-network assignments (Yeo et al., 2011). In addition, we extracted node subsets connecting the previously identified vmPFC regions with bilateral hippocampal and amygdala segmentations (**Figure 1**).

**Figure 1.**
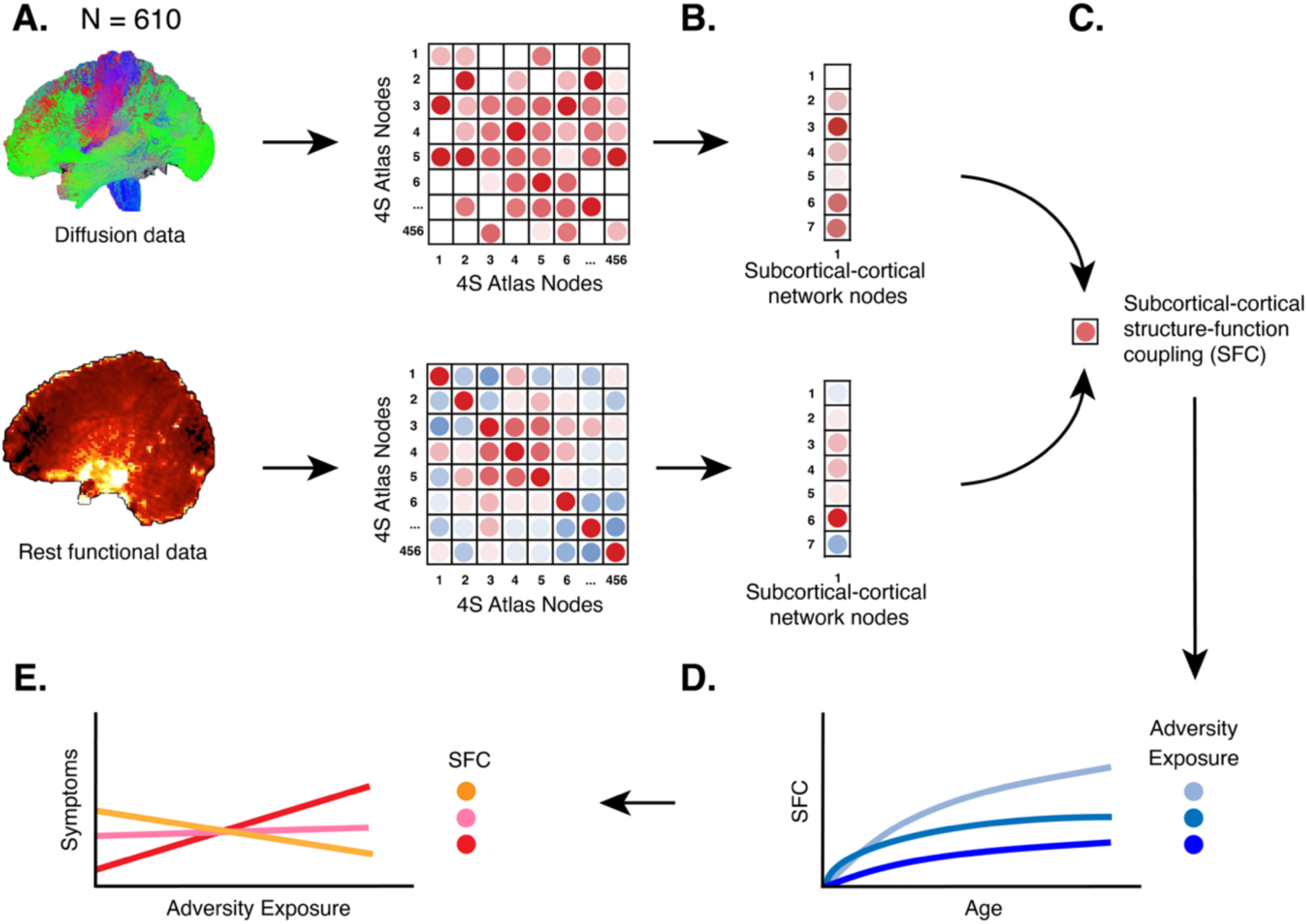
Diagram of analytic workflow. A) Structural and resting state data were parcellated using the 4S atlas to yield structural and functional connectomes. B) Subsets of regions for each matrix were selected to reflect structural and functional connectivity, respectively, within specific circuits of interest. C) Structure-function coupling between connectivity vectors was computed, and the value reflecting subcortical SFC was retained for further analysis. D) GAMs were used to assess relations between age, adversity exposure, and SFC. E) Robust regression models were used to assess the role of subcortical-cortical SFC in moderating associations between adversity and mental health symptoms.

#### Structure-function coupling

SFC was operationalized as the Spearman’s correlation coefficient between each atlas region’s structural and functional connectivity with every other region (Baum et al., 2020). SFC was computed iteratively for subcortical-cortical network connections. Specifically, for each set of cortical nodes, we created a subset matrix of nodes that comprised a given network (e.g., vmPFC; frontoparietal network) along with a subcortical ROI (i.e., the left or right hippocampus or amygdala). The correlation between the structural and functional vectors representing the subcortical node’s connectivity with other cortical nodes in the subsetted matrix was then computed, excluding the self-correlation and edges with a value of 0. The coupling *r-*value was then retained for each subject. This process was repeated for each connection of interest (e.g., hippocampus-vmPFC, amygdala-visual network, hippocampus-default mode network). Left and right subcortical region SFC *r*-values were averaged together. These steps yielded a final dataset that included 14 subnetoworks, corresponding to amygdala and hippocampal connections to each of the seven Yeo functional networks (Yeo et al., 2011) in addition to the amygdala-vmPFC and hippocampal-vmPFC connections. SFC estimates for each connection were then winsorized across participants to mitigate outlier effects. We chose to winsorize rather than omit outliers in order to preserve sample size, but confirmed that all winsorized estimates were correlated at levels greater than 0.95 with the original estimates. Finally, prior to statistical analyses, we corrected SFC measures for site effects using ComBat harmonization while protecting for age and adversity exposure as covariates of interest (Fortin et al., 2018).

### Statistical analyses

Statistical analyses were conducted in R, version 4.5.1.

#### Development of subcortical-cortical connections

We used the *mgcv* package (S. Wood, 2000) to fit generalized additive models (GAMs) to test for linear and nonlinear age-related effects. GAMs were fitted with a smoothing function applied to the age term, with number of basis functions set to k=3 as in previous developmental work (e.g., Sydnor et al., 2023). This smoothed term generates a spline, or a combination of weighted basis functions capable of flexibly capturing linear and nonlinear effects (Jackson, 2024; Simonsohn, 2024). To minimize risk of overfitting, increasing complexity of smoothed terms within the GAM model was penalized using restricted maximum likelihood. In order to systematically evaluate age effects on subcortical-cortical SFC, we fitted two models per connection: a null model, estimated without the smoothed age term, and a full model that included the smoothed age term. Model fits for the null model GAM and the full model GAM were compared using an ANOVA. The magnitude of the age effect was quantified by calculating the partial R^2^ value between the full and null models. All age-related full and null models controlled for sex assigned at birth, mean whole-brain SFC, average root mean squared displacement during the resting-state scans, mean framewise displacement during the diffusion scan, and diffusion quality score. FDR correction was applied to analyses examining age effects on SFC between subcortical regions and the seven Yeo functional networks to account for the seven models fitted for each subcortical region of interest.

#### Interaction effects between age and adversity exposure in subcortical-cortical connections

Given our a-priori hypotheses regarding interactions between adversity exposure and neurodevelopment, we chose to assess impacts of adversity on SFC using an interaction framework. Using a parallel approach to the above, we systematically evaluated potential interaction effects between age and adversity exposure on subcortical-cortical SFC by fitting two models per connection: a null model, which included a smoothed age term but did not include an adversity exposure term, and a full model that included a smoothed age by adversity interaction term as well as a smoothed age term and a smoothed adversity term. The smoothed interaction term between age and adversity exposure was fitted using a tensor product interaction in order to allow for the possibility of both interactive and main effects (Wood, 2017). The null and full models were compared using an ANOVA, and if including the interaction term significantly improved model fit, full model statistics were examined. The magnitude of the interaction term effect was quantified by calculating the partial R^2^ value between the full and null models. Significant interactions were probed using the Johnson-Neyman procedure (Johnson & Fay, 1950). All age-related full and null models controlled for sex assigned at birth, mean whole-brain SFC, average root mean squared displacement during the resting-state scans, mean framewise displacement during the diffusion scan, diffusion quality score, total household income (Noble et al., 2015; Tooley et al., 2021), and education level of the parent participating in the study (Noble et al., 2013). We opted to include proxy measures of socioeconomic status (e.g., parental education, combined family income) as covariates in order to estimate the effects of adversity over and above socioeconomic status-related environmental factors (Amso & Lynn, 2017). FDR correction was applied to analyses examining age effects on SFC between subcortical regions and the seven Yeo functional networks to account for the seven models fitted for each subcortical region of interest.

#### Associations with clinical symptoms

Finally, we sought to determine whether hippocampal-vmPFC and amygdala-vmPFC connections that were associated with adversity exposure moderated associations between adversity and psychopathology. We examined these associations using robust regression, an approach that mitigates the influence of outliers on effect estimations (Todorov & Filzmoser, 2009). Specifically, we fit two robust regression models using the *robustbase* R package (Maechler et al., 2025) for each of the three symptom measures: a null model, which included a linear term for adversity exposure but did not include SFC, and a full model that included an interaction between adversity exposure and SFC as well as main effects of adversity exposure and SFC. The null and full models were compared using an ANOVA, and if including the interaction term significantly improved model fit, full model statistics were examined. The magnitude of the interaction term effect was quantified by calculating the partial R^2^ value between the full and null models. Subsequently, significant interactions were probed using the Johnson-Neyman procedure (Johnson & Fay, 1950). All models included age, sex assigned at birth, primary parent education level, total household income, and primary diagnostic category as covariates. We chose to include primary diagnostic category as a covariate as the HBN sample comprises considerable clinical heterogeneity, including youth diagnosed with conduct disorders, mood disorders, and neurodevelopmental disorders. In particular, a high proportion of the present sample (63%) was diagnosed with a neurodevelopmental disorder, and evidence suggests that the CBCL scale functions differently in youth with neurodevelopmental disorders such as autism (e.g., de la Roche et al., 2024; Volk et al., 2025). Thus, we aimed to mitigate potential differences across disorders by including primary diagnostic category as a covariate in all clinical models.

#### Data and Code Availability

Preprocessed data used in the present study were obtained through open access repositories (Richie-Halford et al., 2022; Shafiei et al., 2025). Code used to compute the structure-coupling measures used in this study, as well as analysis code, can be found at https://github.com/Yale-CANDLab/Sisk_HBN_StructureFunction.

## Results

### Hippocampal-vmPFC and amygdala-vmPFC development

#### Development of subcortical-vmPFC connections

First, we sought to examine the effect of age on SFC in hippocampal-vmPFC and amygdala-vmPFC connections. Inclusion of a smoothed age term did not significantly improve model fit for either amygdala-vmPFC SFC (*p* = .593, partial R^2^ < 0.001) or hippocampal-vmPFC SFC (*p* = .503, partial R^2^ < 0.001), suggesting that the degree to which subcortical-vmPFC functional connectivity is constrained by structural wiring remains largely stable across childhood and adolescence.

#### Interactions between age and adversity exposure in subcortical-vmPFC connections

We next examined whether there were significant interactive effects between age and adversity exposure that were associated with amygdala-vmPFC SFC and hippocampal-vmPFC. Including a smoothed age by adversity exposure interaction term significantly improved model fit for amygdala-vmPFC SFC (*p* = .038, partial R^2^ = 0.015). Further examination of this model revealed that the smoothed interaction between age and adversity was statistically significant (F(1.000) = 7.228, e.d.f. = 1.000, *p* = .007). Neither the smoothed age term (F(1.000) = 0.350, e.d.f. = 1.000, *p* = .554) or the smoothed adversity term (F(1.878) = 0.825, e.d.f. = 1.651, *p* = .353) were independently significant (**Figure 2**). As the effective degrees of freedom for the interaction term suggested a linear effect, we used a linear Johnson-Neyman test to further probe the interaction. Results revealed that for youth with 10 or more adverse exposures, there was a positive association between age and adversity exposure (JN = [-0.02, 9.99]), whereas for individuals with fewer than 10 adverse exposures, the association between age and adversity was not significant. Including a smoothed age by adversity exposure interaction term did not significantly improve model fit for hippocampal-vmPFC SFC (*p* = .314, partial R^2^ = 0.006). These findings suggest that adversity exposure may alter the development of amygdala-vmPFC circuits in youth exposed to high levels of adversity.

**Figure 2.**
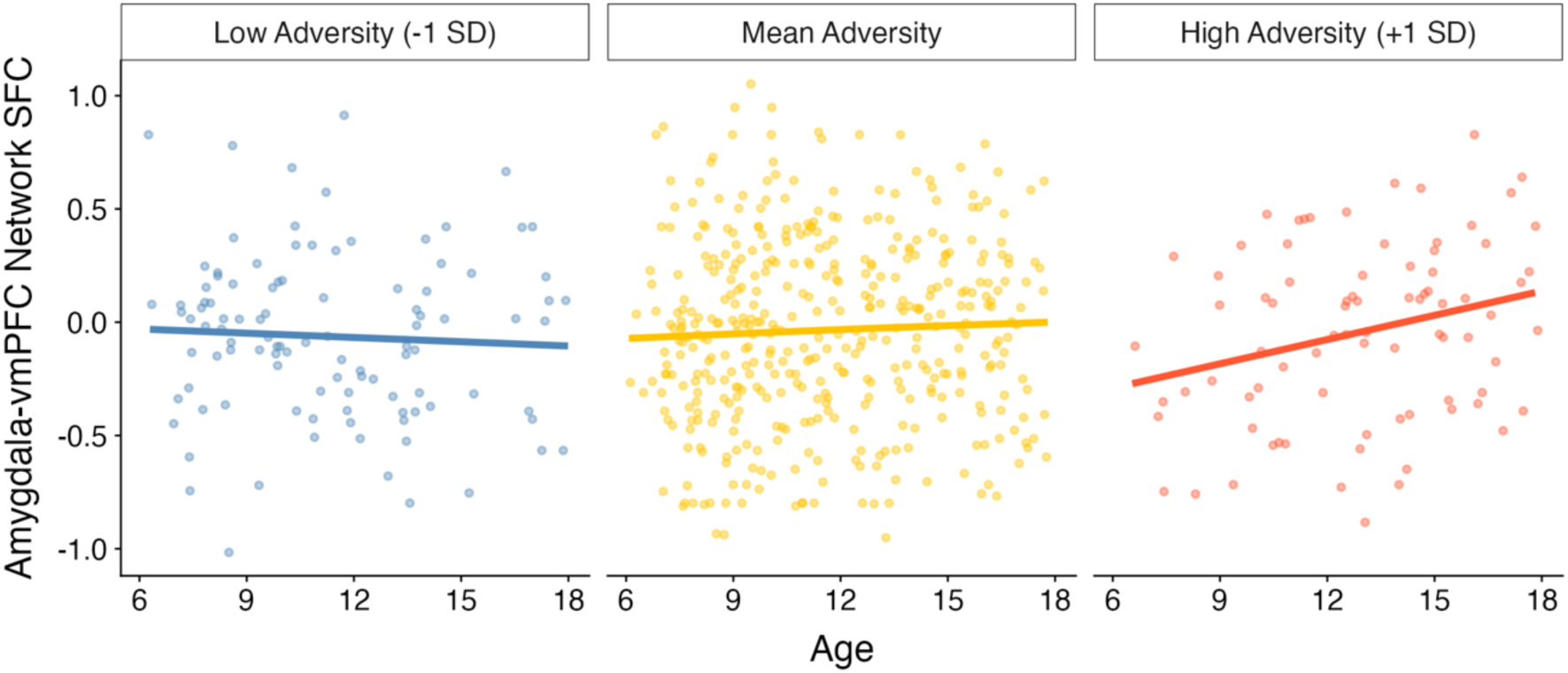
Amygdala-vmPFC SFC is associated with an interaction between age and adversity exposure, such that age is positively associated with adversity exposure for youth with 10 or more (but not fewer than 10) adverse exposures.

#### Associations with clinical symptoms

We next tested whether SFC of amygdala-vmPFC and hippocampal-vmPFC circuits (**Figure 3a**) moderated associations between adversity exposure and clinical symptoms. Indeed, for internalizing symptoms, including an interaction between amygdala-vmPFC SFC and adversity exposure significantly improved model fit relative to the model that included adversity exposure alone (*p =* .018, partial R^2^ = 0.015). The full model showed a significant interaction between adversity exposure and amygdala-vmPFC SFC, such that stronger amygdala-vmPFC SFC was associated with decreased internalizing symptoms for individuals with higher adversity exposure, but not those with lower exposure (β = -0.102, SE = 0.041, *t* = -2.521, *p* = .012; **Figure 3b**). We next probed this interaction using a Johnson-Neyman test (Johnson & Fay, 1950) to identify the significant intervals. Results revealed that the association between adversity exposure and internalizing symptoms was significant for individuals with amygdala-vmPFC SFC less than 0.400 (JN = [0.400, 4.310], **Figure 3c**), whereas for individuals with amygdala-vmPFC SFC greater than 0.400, SFC buffered the association between adversity exposure and internalizing symptoms such that it was no longer significant. Including an interaction between amygdala-vmPFC SFC and adversity exposure did not significantly improve model fit for externalizing symptoms *(p* = .318, partial R^2^ = 0.004) or total symptoms models *(p* = .225, partial R^2^ = 0.005).

**Figure 3.**
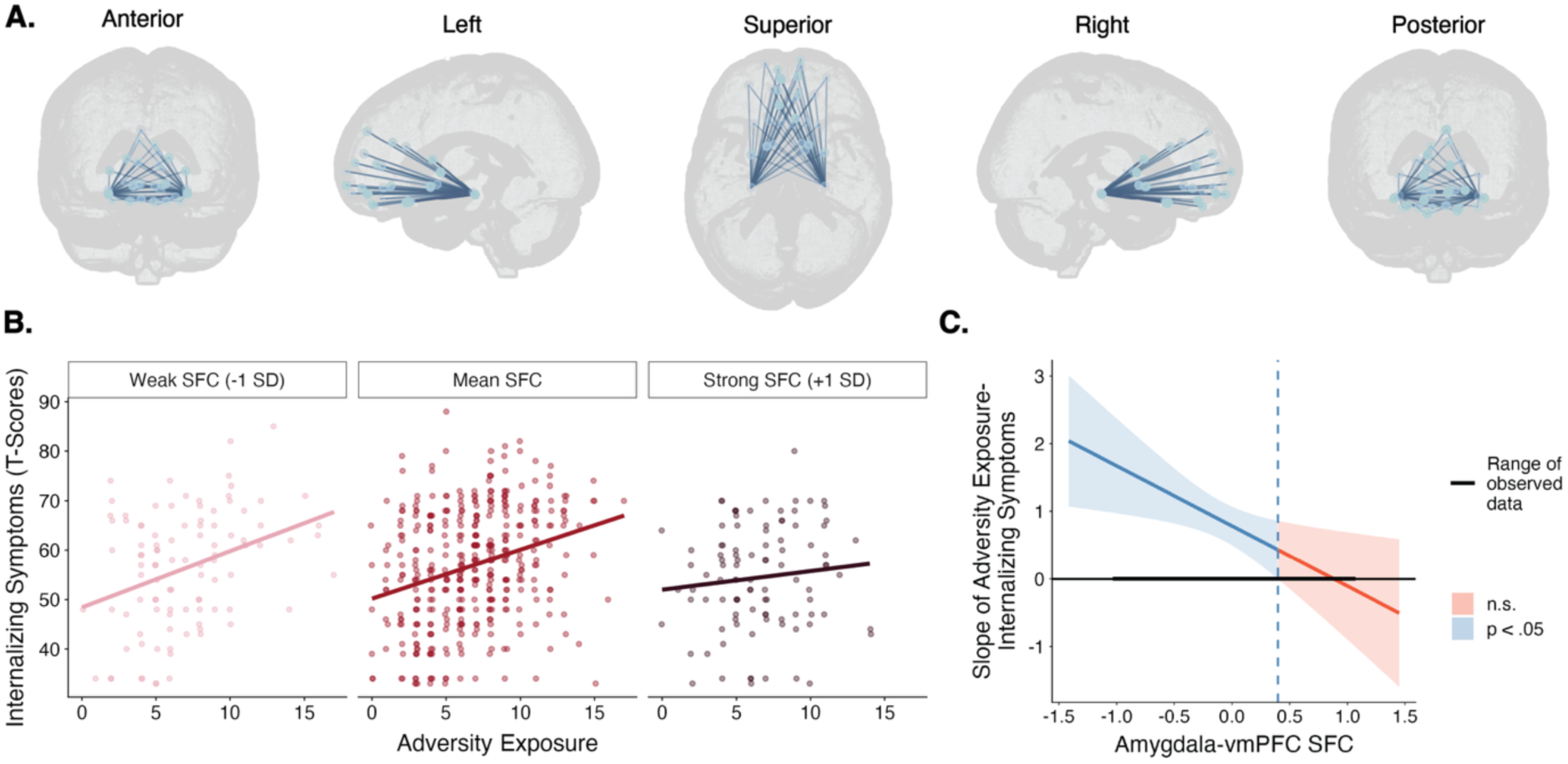
Amygdala-vmPFC SFC moderates the association between adversity exposure and internalizing symptoms. (A) Connections between bilateral amygdalae and nodes included in the vmPFC network. (B) The interaction between amygdala-vmPFC SFC and adversity exposure was associated with internalizing symptoms. Associations between adversity exposure and internalizing syptoms were stratified across three levels of SFC strength (for visualization purposes). (C) A Johnson-Neyman plot depicts the statistically significant interval of the interaction. Amygala-vmPFC SFC significantly moderated the association between adversity exposure and internalizing symptoms, such that adversity exposure remained significantly associated with internalizing symptoms when amygdala-vmPFC SFC was less than 0.400. When SFC was above 0.400, amygdala-vmPFC SFC buffered the association such that adversity exposure was no longer significantly associated with internalizing symptoms.

**Figure 4.**
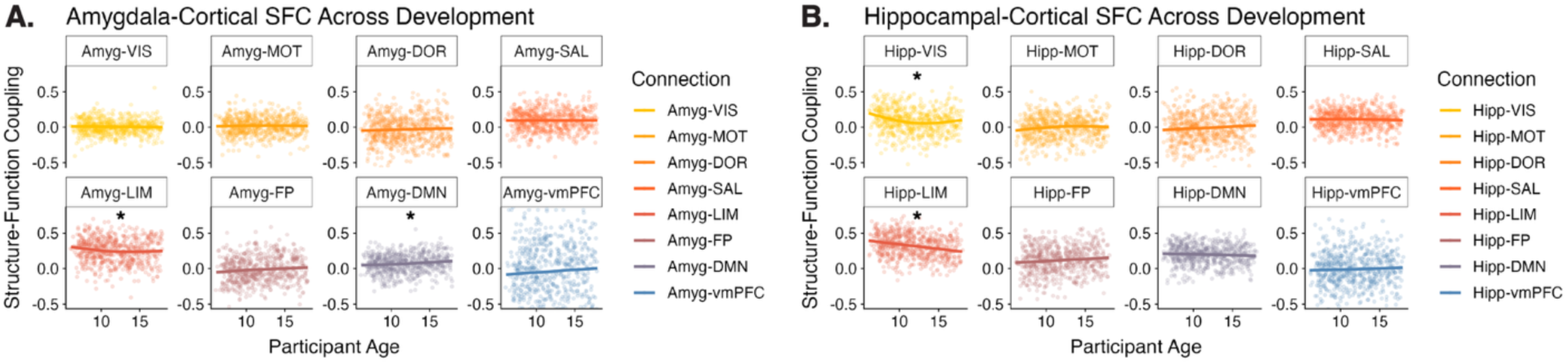
Associations between age and SFC among A) amygdala-cortical and B) hippocampal-cortical connections. Significant age effects are denoted with an asterisk. DMN = default mode network, FP = frontoparietal network, LIM = limbic network, SAL = salience network, DOR = dorsal attention network, MOT = somatomotor network, VIS = visual network, vmPFC = ventromedial prefrontal cortex.

With regard to hippocampal-vmPFC SFC, we found that including an interaction between hippocampal-vmPFC SFC and adversity exposure did not significantly improve model fit for internalizing symptoms *(p* = .159, partial R^2^ = 0.006), but did significantly improve model fit for total symptoms (*p =* .048, partial R^2^ = 0.008), and marginally improved model fit for externalizing symptoms (*p =* .051, partial R^2^ = 0.009). Further examination of the full model for total symptoms revealed a significant interaction between adversity exposure and hippocampal-vmPFC SFC (β = -0.084, SE = 0.040, *t* = -2.082, *p* = .038). Examination of this interaction using the Johnson-Neyman procedure revealed that stronger hippocampal-vmPFC SFC buffered the association between adversity exposure and total symptoms. Specifically, the association between adversity exposure and total symptoms was significant for individuals with hippocampal-vmPFC SFC less than 0.340 (JN = [0.330, 7.670]), whereas for individuals with hippocampal-vmPFC SFC greater than 0.340, SFC buffered the association between adversity exposure and internalizing symptoms such that it was no longer significant. Together, these results suggest that among youth with higher levels of adversity exposure, greater SFC in the amygdala-vmPFC circuit may buffer specifically against higher internalizing symptomatology, whereas greater SFC in the hippocampal-vmPFC circuit may buffer against higher total symptoms.

### Hippocampal-functional network and amygdala-functional network development

#### Development of subcortical-functional network connections

Next, we sought to characterize age-related change among SFC connections between the amygdala, hippocampus, and the seven Yeo functional networks (Yeo et al., 2011). Inclusion of a smoothed age term resulted in significantly improved model fit for amygdala-limbic network SFC (*p =* 0.012, partial R^2^ = 0.015), amygdala-default mode network SFC (partial R^2^ = 0.007, *p* = .041), hippocampal-limbic network SFC (*p <* .001, partial R^2^ = 0.046), and hippocampal-visual network SFC (*p <* .001, partial R^2^ = 0.040). Further examination of these models revealed a negative effect of age on amygdala-limbic SFC (F(1.950) = 4.833, e.d.f. = 1.777, *p* = .020), amygdala-default mode SFC (F(1.001) = 4.159, e.d.f. = 1.000, *p* = .042), hippocampal-limbic SFC (F(1.001) = 28.700, e.d.f. = 1.000, *p* < .001), and hippocampal-visual SFC (F(1.996) = 11.390, e.d.f. = 1.938, *p* < .001). Age effects for hippocampal-limbic and hippocampal-visual SFC survived FDR correction across the 7 network models per subcortical region (*ps_FDR_* < .001).

#### Interactions between age and adversity exposure in subcortical-functional network connections

In order to test the extent to which the age by adversity interaction was specifically associated with SFC in the amygdala-vmPFC circuit, we next examined whether the interaction between age and adversity exposure was associated with SFC in connections between the hippocampus, amygdala, and the seven Yeo functional networks. Inclusion of a smoothed age by adversity exposure interaction term significantly improved model fit for hippocampal-limbic SFC alone (*p =* .028, partial R^2^ = 0.019). Further examination revealed that the improved model fit was driven by separate main effects of age and adversity exposure. Refitting the model with main effects (excluding the nonsignificant age by adversity interaction) yielded significant main effects of age (F(1.000) = 34.227, e.d.f. = 1.000, *p* < .001) and adversity exposure (F(1.953) = 2.933, e.d.f. = 1.784, *p* = .037). However, the improvement in model fit afforded by inclusion of the adversity by age interaction was not significant after correcting for the seven models fitted for each of the Yeo functional networks (*p_FDR_ =* .196).

#### Exploratory analyses

Given that hippocampal-limbic SFC showed a main effect of adversity exposure, we examined whether SFC in this circuit moderated associations between adversity exposure and clinical symptoms. With regard to internalizing symptoms, including an interaction between hippocampal-limbic SFC and adversity exposure significantly improved model fit relative to the model that included adversity exposure alone (*p =* .034, partial R^2^ = 0.005). The full model showed a significant interaction between adversity exposure and hippocampal-limbic SFC for internalizing symptoms, such that weaker hippocampal-limbic SFC was associated with decreased symptoms for individuals with higher adversity exposure, but not lower adversity exposure (β = 0.098, SE = 0.038, *t* = 2.567, *p =* .010). We probed this interaction using the Johnson-Neyman procedure, which revealed that the interaction between adversity exposure and hippocampal-limbic SFC was significant for individuals with hippocampal-limbic SFC greater than 0.120 (JN = [-1.470, 0.120]) (**Figure 5**), whereas for individuals with hippocampal-limbic SFC less than 0.120 the association between adversity exposure and internalizing symptoms was no longer significant. With regard to total symptoms, including an interaction between hippocampal-limbic SFC and adversity exposure marginally improved model fit relative to the null model (*p =* .057, partial R^2^ = 0.002), and did not significantly improve model fit for externalizing symptoms *(p* = .153, partial R^2^ = 0.004).

**Figure 5.**
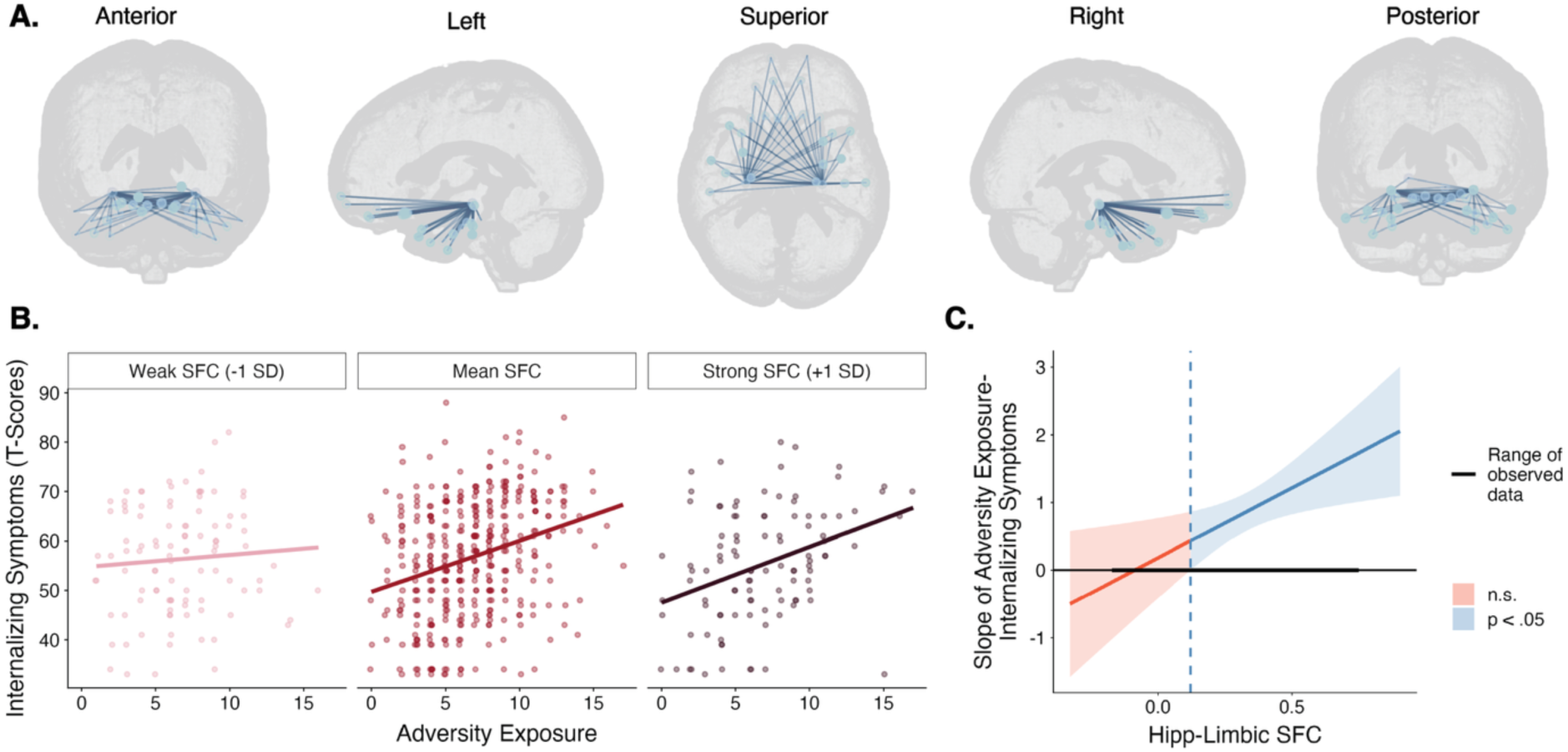
Hippocampal-limbic network SFC moderated the association between adversity exposure and internalizing symptoms. A) Connections between bilateral hippocampi and nodes comprising the limbic functional network. B) The interaction between adversity exposure and hippocampal-limbic network SFC was associated with internalizing symptoms. Here, for visualization purposes, associations between adversity exposure and internalizing symptoms were stratified across three levels of SFC strength (for visualization purposes). C) A Johnson-Neyman plot depicts the significant interval of the interaction. Specifically, adversity exposure was significantly associated with internalizing symptoms when hippocampal-limbic SFC was greater than 0.090. When SFC was less than 0.09, SFC buffered the association such that adversity exposure was no longer significantly associated with internalizing symptoms.

## Discussion

Variability in the structural and functional development of bidirectional connections linking the hippocampus and amygdala with the prefrontal cortex underlies individual differences in stress responding, cognition, and mental health (Casey et al., 2019). In the present study, we demonstrated that adversity exposure alters the coupling of structural and functional connectivity of amygdala-prefrontal and hippocampal-limbic circuits, and that altered SFC buffers the effects of adversity exposure on mental health. We identified adversity-related differences in the development of amygdala-vmPFC SFC: whereas youth with lower levels of adversity exposure showed relatively stable amygdala-vmPFC SFC with age, youth exposed to higher levels of adversity showed increased amygdala-vmPFC SFC with age. Furthermore, among youth with higher levels of adversity exposure, the association between adversity exposure and internalizing symptoms was attenuated for those with strong amygdala-vmPFC SFC. In parallel, hippocampal-limbic network SFC was negatively associated with adversity exposure across development and weaker hippocampal-limbic SFC attenuated the association between adversity exposure and internalizing symptoms for youth with higher adversity exposure. Importantly, these patterns were circuit-specific, and were not observed in other amygdala- or hippocampal-cortical connections. Together, these findings suggest that amygdala-prefrontal and hippocampal-limbic network SFC may capture adversity-related circuit remodeling during development, and that adaptive patterns of SFC may differ between stress regulation and memory circuits in ways that moderate individual-level risk for or resilience against internalizing psychopathology.

Bidirectional projections between the amygdala and vmPFC undergo dynamic changes over the course of development (Gabard-Durnam et al., 2014; Ojha et al., 2025) and may develop differently in youth exposed to adversity (Brieant et al., 2021; Gee et al., 2013; Herzberg et al., 2021). In rodents, neuronal projections from the mPFC to the amygdala undergo significant maturation during the juvenile and early adolescent stages, and increased inhibitory neurotransmission during adolescence suggests that this stage may represent a sensitive period for amygdala-prefrontal circuit development (Arruda-Carvalho et al., 2017). In the present study, we observed an age-related increase in amygdala-vmPFC SFC among youth with higher levels of adversity exposure. As SFC is sensitive to myelination and the neuronal excitation-to-inhibition balance, both putative mechanisms of plasticity restriction (Fotiadis et al., 2023), stronger amygdala-vmPFC SFC in youth exposed to more adversity may reflect experience-dependent increases in plasticity restriction factors in this circuit. Additionally, stronger inhibitory connectivity within the amygdala-vmPFC pathway may indicate more effective regulation of amygdala reactivity by the prefrontal cortex (e.g., He et al., 2023; Tottenham & Gabard-Durnam, 2017). In this context, increased amygdala-vmPFC SFC in youth with higher adversity exposure may represent ontogenetic adaptation to facilitate improved downregulation of the neural stress response. In line with our hypotheses, amygdala-vmPFC SFC in youth with higher levels of adversity exposure buffered the association between adversity exposure and internalizing symptoms, suggesting that adversity-related increases in amygdala-vmPFC SFC may facilitate resilience against psychopathology.

Paralleling amygdala-vmPFC circuit development, hippocampal-cortical circuits also undergo protracted development that continues into adolescence (Calabro et al., 2020; Casey et al., 2019; Cruz-Sanchez et al., 2025). We observed a developmental decrease in SFC between the hippocampus and regions comprising the limbic cortical network, primarily located in entorhinal, parahippocampal, and orbitofrontal cortical regions. Prior studies have observed low SFC in cortical regions within the limbic network (Baum et al., 2020; Fotiadis et al., 2023), and we posit that age-related decreases in hippocampal-limbic network SFC may reflect increasing functional signal diversity and flexibility within the hippocampal-limbic circuit over the course of development. The present study further found that SFC in this circuit was negatively associated with adversity exposure, such that higher levels of exposure to adversity were associated with weaker SFC. This pattern may represent adversity-related accelerated neurodevelopment within this circuit. As hippocampal-limbic network connectivity is involved in core cognitive functions such as memory encoding, retrieval, and recall (Chao et al., 2020; Raud et al., 2023; Setton et al., 2022), we posit that adversity-related changes in the development of SFC within the hippocampal-limbic circuit may be linked with earlier emergence of more mature cognitive functioning, potentially alongside adversity-related alterations in learning and memory (Hardi et al., 2025; Pattwell & Bath, 2017). Finally, SFC within the hippocampal-limbic network circuit moderated the association between adversity exposure and internalizing symptoms among youth with higher levels of adversity exposure, such that weaker hippocampal-limbic network SFC attenuated the association between adversity exposure and internalizing symptoms. Together, these findings suggest a distinct pattern of adversity-related neural adaptation within the hippocampal-limbic circuit.

The opposing patterns of SFC changes observed in amygdala-vmPFC and hippocampal-limbic circuits among adversity-exposed youth suggest that SFC varies adaptively by circuit. That is, our findings suggest that both *weaker* SFC in hippocampal-limbic network connections and *stronger* SFC in amygdala-vmPFC connections moderate the association between adversity exposure and internalizing symptoms. These distinct patterns of SFC may reflect differential circuit functions. Strengthened inhibitory connectivity between the amgydala and vmPFC may facilitate prefrontal regulation of amygdala reactivity, which has been linked with more resilient functioning (Sisk et al., 2025; van Rooij et al., 2024; Zhang et al., 2023). Weaker SFC in hippocampal-limbic network connections may reflect increased functional signal diversity (Fotiadis et al., 2023), as well as earlier emergence of a more mature circuit phenotype that supports more adult-like cognition (Bath et al., 2016; Callaghan & Tottenham, 2016). Future studies that jointly examine behavioral indicators of cognitive performance and stress responding together with multimodal neuroimaging data would be well-positioned to further examine this hypothesis.

The present study has a number of strengths, including a relatively large sample of youth, rigorous characterization of multimodal links between brain structure and function, and robust processing and analytic methods. Furthermore, the sample included individuals with a wide range of clinical diagnoses, as well as a wide range of adversity exposure, capturing an important cross-section of youth experience that may result in findings that are more likely to generalize (Marek & Laumann, 2024). However, there are several important limitations to note. While we used the total exposure score from the Negative Life Events Scale (Tiet et al., 2001) to model adversity exposure, accounting for additional features of adverse events and the contexts in which they occur can help to explain variability in outcomes (Cohodes et al., 2021; Ellis et al., 2022; Gee & Casey, 2015; McLaughlin et al., 2014; Pollak & Smith, 2021). In future work, accounting for features such as developmental timing of adversity (Cohodes et al., 2023; Pechtel et al., 2014; Sicorello et al., 2021; Sisk, Keding, Cohodes, et al., 2025; Sisk, Keding, Ruiz, et al., 2025; Zhu et al., 2023) may yield greater insight into the precise temporal dynamics of neurodevelopment following adversity exposure. We also controlled for differences in maternal education and total household income in the present study. While this approach facilitates isolation of the effects of adversity relative to the child’s larger family-level environmental context, adversity exposure and indices of socioeconomic status are often intertwined (Mills-Koonce & Towe-Goodman, 2012); this approach may thus dilute the effect size observed for adversity exposure. It is also important to note that inter-regional tractography, as was used to construct the structural connectome, has several known limitations. For instance, streamlines exhibit a relatively high false positive rate and distance-dependent bias (e.g., Maier-Hein et al., 2017; Reveley et al., 2015). While we attempted to mitigate these limitations by using a weighted thresholding approach, future advances in estimating correspondence between neural structure and function may improve upon the present methods. Finally, this study is limited to cross-sectional inference into the development of subcortical-cortical SFC. Leveraging data from large-scale longitudinal studies such as the Adolescent Brain Cognitive Development (Casey et al., 2018) and Healthy Brain and Child Development (Volkow et al., 2024) studies to investigate how these patterns emerge within individuals will provide valuable insight into the extent to which the present findings are recapitulated within-person.

### Conclusions

The present study identified adversity-related disruption in coupling between the structure and function of amygdala-vmPFC and hippocampal-limbic circuits across development. Youth with higher adversity exposure showed greater age-related increases in amygdala-vmPFC SFC, relative to youth with lower exposure, potentially reflecting adaptive neurodevelopmental change. Adversity exposure was also negatively associated with hippocampal-limbic SFC across development. Among youth with higher (rather than lower) adversity exposure, strong amygdala-vmPFC SFC and weak hippocampal-limbic SFC buffered the association between adversity exposure and internalizing symptoms. Taken together, these findings suggest that adversity exposure jointly shapes developing brain structure and function in a manner that may promote resilience against worsened internalizing symptomaology. Evidence of differential effects of adversity exposure on SFC—stronger amygdala-vmPFC SFC and weaker hippocampal-limbic SFC—points to circuit-specific adaptations that contribute to individual variation in risk and resilience.

## Acknowledgements

We would like to thank the participants and data collection teams involved in the Healthy Brain Network study, as well the research teams whose efforts to preprocess the HBN neuroimaging data and release the derivatives publicly made this study possible. This manuscript was prepared using a limited access dataset obtained from the Child Mind Institute Biobank (Healthy Brain Network study). This manuscript reflects the views of the authors and does not necessarily reflect the opinions or views of the Child Mind Institute.

